# Flavobacteria buffer nitrous oxide emissions from partial denitrifiers in coastal sediments

**DOI:** 10.1101/2025.11.24.689969

**Authors:** Thanh Nguyen-Dinh, Tess Hutchinson, Francesco Ricci, Hanif Prayitno, Luis Jimenez, Vera Eate, Pok Man Leung, Rachael Lappan, Sukhwan Yoon, Wei Wen Wong, Perran L.M. Cook, Chris Greening

## Abstract

Nearly one-fifth of global emissions of the potent greenhouse gas nitrous oxide (N_2_O) originate from the ocean, particularly from nutrient-polluted coastal regions. Permeable (sandy) sediments, which cover half of the continental shelf worldwide, are potential sources of N_2_O due to increasing nutrient inputs from urbanization and agriculture. Yet, the microbial processes determining N_2_O emissions in these dynamic and unique ecosystems remain understudied. Here, we combined environmental measurements, bacterial cultivation, and genomic analyses to understand the microbes and processes controlling N_2_O cycling in permeable sediments from Port Phillip Bay (Australia). We established a genomic resource comprising 249 metagenome-assembled genomes and 95 new isolate genomes. Genome-based metabolic reconstructions and culture-based gas measurements revealed diverse bacteria in these sediments produce N_2_O through incomplete denitrification pathways. However, these bacteria co-occurred with highly abundant clade II N_2_O-reducing bacteria from the *Flavobacteriaceae* family. Kinetic profiling revealed both clade II *nosZ* flavobacterial isolates and whole sand communities exhibit a low affinity for N_2_O, contrary to previous reports that clade II N_2_O reducers generally have a high substrate affinity. This indicates adaptation to the high residence times of N_2_O within production and consumption zones in the sands. Collectively, these N_2_O reducers remove most N_2_O produced in permeable sediments, supporting lower-than-expected coastal emissions predicted by biogeochemical models. We conclude that permeable sediments host specialised microbial communities that mitigate N_2_O emissions and buffer marine nitrogen cycling amid rising nutrient pollution.

## Introduction

Nitrous oxide (N_2_O) is a long-lived atmospheric trace gas contributing to stratospheric ozone depletion and climate change [1]. With a warming potential approximately 300 times higher than that of carbon dioxide on a 100-year time scale, N_2_O is the third largest contributor (6.4%) to total greenhouse gas emissions after CO_2_ and CH_4_ [2,3]. Up to 57% of the total N_2_O emissions come from natural sources, in which marine ecosystems contribute to nearly half of such emissions at 2-7 Tg N year^−1^ [4]. Estuarine and coastal zones function as a natural biofilter, removing excess nutrients at the interface between land and ocean [5]. However, increasing anthropogenic activities, such as nutrient loading, aquaculture and fertiliser use, are inputting nitrogen to these ecosystems, which in turn results in widespread eutrophication and exacerbates N_2_O emissions [4,6–9]. N_2_O emissions from coastal environments remain insufficiently understood and poorly quantified in global N_2_O budgets due to complex N_2_O production pathways and high spatiotemporal variability [7,10,11].

Permeable sediments (i.e. sands and gravels) dominate coastal regions, covering nearly half of the continental shelf worldwide [12]. These sediments are subject to highly turbulent hydrodynamic conditions that vary across space and time, as opposed to fine-grained impermeable (cohesive) sediments [13–15]. Advective pore-water flows drive continuous exchange of dissolved solids and gases, suspended particles and microorganisms between the sediment and overlying water column, making redox conditions in the sediments highly variable over short distances and timescales [13–15]. With microbial abundance of ∼10^9^ cells cm^−3^ sediments [16], sands affect N_2_O emissions in coastal regions due to enhanced microbial nitrogen cycling linked to anthropogenic nitrogen inputs. Nitrification and denitrification are the two main microbial processes that control the N_2_O budget in aquatic environments [17,18]. In well-oxygenated waters, N_2_O can be released by ammonia-oxidizing bacteria (AOB) and archaea (AOA) as a byproduct of ammonia oxidation to nitrite (NO_2_ ^−^) (i.e. the first step of nitrification) [9,18]. By contrast, N_2_O is produced as an intermediate during denitrification of nitrate (NO_3_ ^−^) or NO_2_ ^−^ under oxygen-depleted conditions [17,19]. Many denitrifiers lack the genetic capacity for N_2_O reduction, resulting in N_2_O released as the final product instead of inert dinitrogen gas (N_2_) [20,21]. Furthermore, abiotic processes, such as (photo)-chemodenitrification, can also contribute to the production of N_2_O in marine waters and sediments [22,23].

Despite several processes leading to N_2_O production, the major biological sink of this gas is its reduction to dinitrogen gas (N_2_) by microorganisms expressing the periplasmic, copper-containing enzyme N_2_O reductase (NosZ) [24,25]. Phylogenetic analyses resolve the enzyme into clade I (NosZI), clade II (NosZII) and the recently defined clade III (NosZIII) [26–28]. NosZI is often associated with complete denitrifiers. In contrast, NosZII is found in much more diverse microbial taxa that typically lack key denitrification enzymes (i.e. NirK and NirS), suggesting N_2_O serves as an electron acceptor even for non-denitrifiers [27–29]; for example, some non-denitrifying bacteria encoding the NosZII, including the acidophilic bacterium *Desulfosporosinus nitrosoreducens* [30] and methanotrophs *Methylocella tundrae* and *Methylacidiphilum caldifontis* [31], reduce N_2_O to conserve energy through anaerobic respiration, while others use it to dispose excess electrons during transient anoxia (Park et al., 2017). Beyond these two clades, a recent study discovered a widespread and active clade III lactonase-type NosZ that is divergent in sequence but similar in structure to NosZI and NosZII [26]. These findings have expanded our understanding on the diversity of N_2_O reducers and provide opportunities for mitigating N_2_O emissions.

Given the global extent of permeable sediments, any imbalances between N_2_O production and consumption can greatly impact marine N_2_O budgets. While there is evidence that these sediments are a potential source of N_2_O [32–36], recent hydrodynamic-biogeochemical models suggest that their contribution to coastal N_2_O emissions are relatively minor [37], highlighting uncertainties in the N_2_O budget of these sediments. Furthermore, current understanding of the microbial basis of N_2_O cycling within permeable sediments is largely limited to community surveys [33], with little functional and ecophysiological characterization. In this study, we addressed these knowledge gaps by investigating the functional diversity and capacity of N_2_O-cycling microorganisms in sandy sediments from moderate to high nutrient-impacted beaches in Port Phillip Bay, Australia. To do so, we integrated bacterial cultivation, metagenomic analyses, and biogeochemical assays to provide a newly compiled genome collection of microorganisms inhabiting permeable sediments and assess their role in N_2_O cycling. Altogether, we demonstrate that diverse N_2_O producers and consumers are present and active in coastal sediments, with their tight coupling buffering the N_2_O emissions from these sediments. These findings provide microbial evidence supporting hydrodynamic models [37] that coastal N_2_O emissions may be overestimated and reveal new mediators of nitrogen and greenhouse gas cycling in marine ecosystems.

## Results

### Microbial communities in permeable sediments rapidly consume high concentrations of N_2_O

To determine whether N_2_O production occurs *in situ,* we measured N_2_O in pore waters of subtidal sandy sediments at St. Kilda and Werribee beaches located in Port Phillip Bay, a nitrogen-limited marine embayment in Victoria, Australia. Trace amounts of N_2_O (∼ 0.0055 _µ_M or 84.6% saturation) were detected in some samples; however, N_2_O was generally below the limit of detection (0.004 _µ_M dissolved gas concentration) in porewater across most sampling depths and times (**Fig. S1, Table S9**). The absence of detectable *in situ* N_2_O concentrations is surprising given the denitrification activity previously reported at the same studied sites [38,39]. This may reflect three possibilities (or a combination of): (1) denitrification steps are tightly coupled, producing minimal N_2_O and efficiently reducing it to N_2_, (2) N_2_O is only sporadically produced when NO_3_^−^ is transported into anoxic zones or (3) non-denitrifying bacteria with clade II N_2_O reductases rapidly consume N_2_O produced by other community members.

We performed a series of slurry experiments to determine whether microbial communities within permeable sediments consume N_2_O at a range of concentrations. Under anoxic conditions, microbial communities in the slurries rapidly consumed 0.8 _µ_M and 108 _µ_M N_2_O with the average rates of 0.65 and 49.8 nmol h^−1^ g^−1^ wet sediment, respectively (**Fig. 1A**). Additionally, Werribee slurries showed significantly higher N_2_O consumption rates (*P* = 0.015 -Student’s t-test) than St. Kilda slurries at both concentrations, potentially reflecting long-term enrichment of denitrifying populations in these sediments at the vicinity of a wastewater treatment plant. Kinetic experiments were used to determine whether beach sand microbial communities are adapted to high or low N_2_O concentrations. N_2_O reduction by the sedimentary communities followed Michaelis-Menten-type kinetics (R^2^ = 0.99 for St Kilda, 0.93 for Werribee) with apparent half-saturation constants [K_m(app)_] of 43 and 55 µM N_2_O respectively and maximum reduction rates [V_max (app)_] of 19.6 and 46.3 nmol N_2_O h^−1^ g^−1^ wet sediment (**Fig. 1B**). The [K_m(app)_] values observed in this study are unusually high for marine communities, for example being orders of magnitude higher than those observed in anoxic estuarine water in Chesapeake Bay, US (0.647 µM N_2_O) [25] or those in the oxic-anoxic interface from the eastern Pacific oxygen minimum zone (0.334 µM N_2_O) [40]. This suggests that microbial communities within sands are exposed to much higher N_2_O fluxes compared to the water column, presumably due to nutrient enrichment in these ecosystems. Furthermore, the [V_max(app)_] for N_2_O reduction in these sediments were one order of magnitude higher than previously reported [V_max(app)_] values for denitrification in permeable sediment in Port Phillip Bay (0.5 – 5 nmol g^−1^ h^−1^) [39], implying the capacity for N_2_O consumption exceeded N_2_O production in these sediments. Altogether, our results demonstrate that while permeable sediments could potentially produce elevated N_2_O levels due to nutrient enrichment, rapid microbial consumption helps prevent N_2_O accumulation and emissions. Thus, these sediments may function as hotspots of N_2_O turnover, acting as a dynamic sink despite high N_2_O exposure.

**Figure 1.**
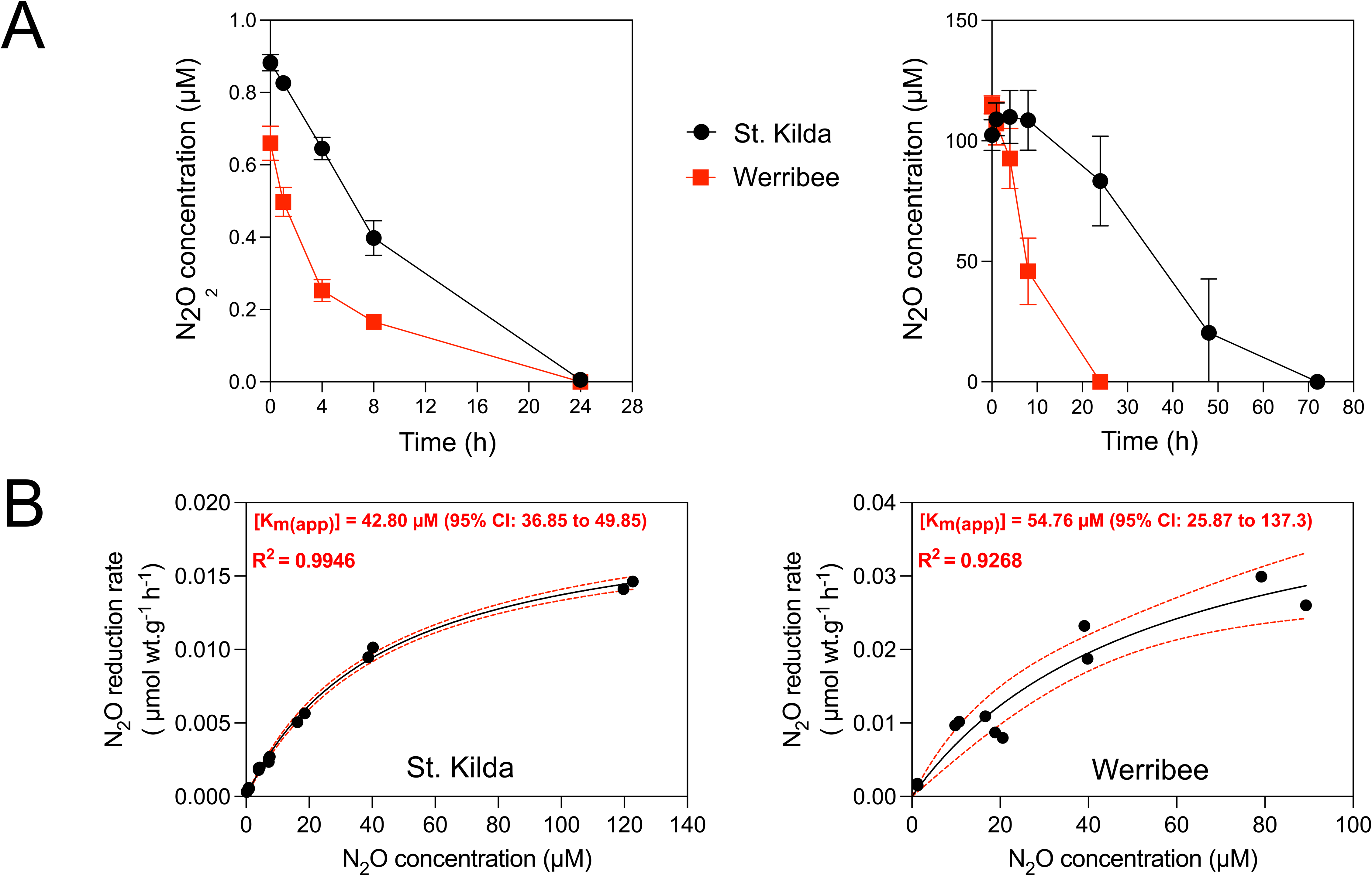
N_2_O consumption by permeable sediments. (**A**) Anoxic slurry incubation of surface sediments (0-5 cm) from St. Kilda (13/05/2024) and Werribee (23/09/2024) with different N_2_O concentrations. (**B**) Kinetics of N_2_O reduction by surface sediments in St. Kilda (13/05/2024 samples) and Werribee (20/08/2025 samples). The curves were fitted and kinetic parameters were approximated based on a Michaelis-Menten non-linear regression model.

### Permeable sediments host diverse N_2_O producers and consumers

We used genome-resolved metagenomics to better understand the microbial and enzymatic mediators of N_2_O cycling in permeable sediments. We sequenced and performed *de novo* co-assembly and binning of metagenomes from the Werribee and St. Kilda beaches, yielding 53 medium-quality and 17 high-quality metagenome-assembled genomes (MAGs). We additionally reanalysed 179 MAGs we previously obtained from St. Kilda permeable sediments [38,41] (**Table S3**). Overall, the resultant 249 MAGs span 19 phyla and 74 orders including the most abundant taxa previously reported for permeable sediment (e.g. *Woeseiaceae*, *Rhodobacteraceae* and *Flavobacteriaceae*) (**Fig. 2B, Table S3**). We then annotated the short reads, derived MAGs and unbinned contigs to determine the functional capacities and diversity of the permeable sediment microbiome.

**Figure 2.**
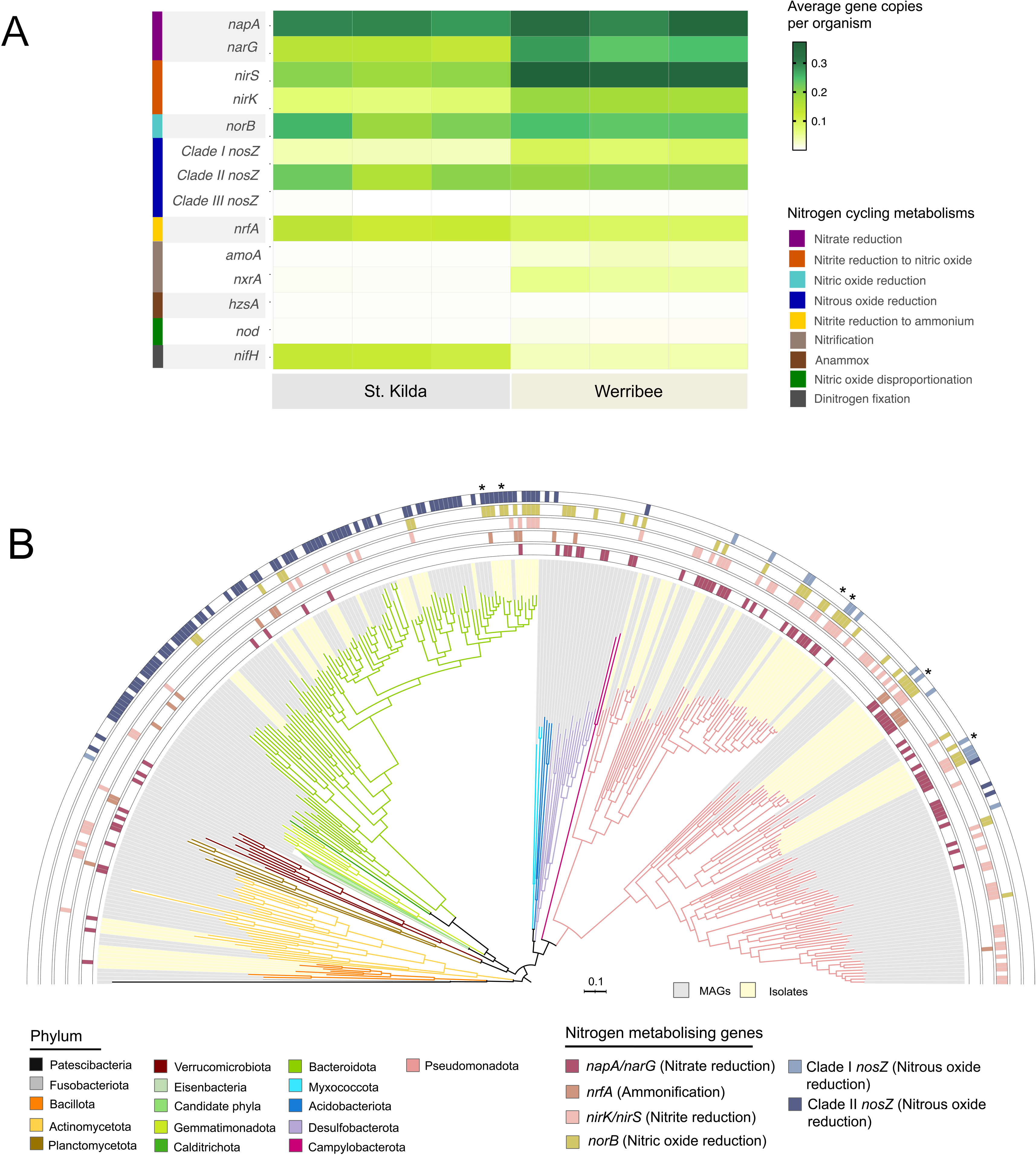
Nitrogen-cycling metabolic capacity of microbial communities in permeable sediments. (**A**) Relative abundance of nitrogen metabolizing genes in St. Kilda and Werribee beaches based on gene-centric metagenomic analysis (**B**) Maximum-likelihood phylogenomic tree showing nitrogen-cycling metabolic gene inventories of 95 bacterial isolates and 245 bacterial MAGs generated from this study and acquired from (Chen et al., 2022, 2021). The tree was built with 1,000 ultrafast bootstrap replicates using the LG+F+I+G4 model via IQ-TREE. Tree branches were colour-indexed to show phylum-level taxonomy of bacterial isolates and MAGs. Key metabolic genes for denitrification were highlighted using coloured rings. Specific isolates were selected for further analysis, including culture-based activity and kinetic measurements (asterisks).

In line with previous studies [38,41], microbial communities within permeable sediments have high capacity for denitrification. Specifically, many community members are predicted to mediate NO_3_^−^ reduction via Nar and Nap (St. Kilda: 44%; Werribee: 58%), NO_2_^−^ reduction via NirS and NirK (St. Kilda: 27%; Werribee: 54%) and nitric oxide (NO) reduction to N_2_O via Nor (St. Kilda: 22%; Werribee: 24%) (**Fig. 2A; Table S2**). Metabolic reconstruction showed that 76 MAGs (30%) encode partial denitrification pathways, i.e. MAGs missing one or more denitrifying reductases (**Table S3**). To minimize artefacts of missing genes due to fragmented genomes, we performed more focused analyses of 64 high-quality MAGs (>90% genome completeness) to more reliably infer partial denitrification pathways. Among these, 16 MAGs were found to possess one or more denitrying reductases but lack the N_2_O reductases. MAGs associated with *Arenicellaceae* (Gammaproteobacteria)*, JABDKV01* (Gammaproteobacteria) and *Pyrinomonadaceae* (Acidobacteriota) families encoded Nor for N_2_O production, suggesting these microorganisms are N_2_O producers within the sand communities. Compared to denitrification, fewer community members mediate nitrification via Amo and Nxr (St. Kilda: 0.4%; Werribee: 8%) or dissimilatory nitrate reduction to ammonium (DNRA via Nrf – St. Kilda: 13%; Werribee: 9%) (**Fig. 2A; Table S2**).

Consistent with our activity measurements, analysis of metagenomic short reads shows that many bacteria encode genes to reduce N_2_O via NosZ (St. Kilda: 23.3%; Werribee: 29.1%; **Fig. 2A**). In detail, bacteria encoding the NosZII are more abundant than the NosZI (19.6% and 20.2% versus 3.5% and 8.7% of St. Kilda and Werribee communities, respectively) while only a minor fraction of the community (St. Kilda: 0.01%; Werribee: 0.8%) encoded the NosZIII (**Fig. 2A**). Consistently, at the MAG level, only five MAGs associated with Pseudomonadota (n = 4) and Verrucomicrobiota (n = 1) encode NosZI whereas 61 MAGs spanning 8 phyla encode the NosZII (**Fig. 2B**). Of NosZII containing MAGs, 49 (80%) were Bacteroidota and 57% of these were *Flavobacteriaceae* family. Furthermore, the majority of NosZII harbouring MAGs (79%) contained only *nosZ* but not any other denitrifying genes, suggesting these microbial groups only consume, but do not produce N_2_O.

To gain further insight into the phylogenetic diversity of the N_2_O-reducers, we constructed a phylogenetic tree of NosZ including 647 amino sequences retrieved from 36 isolate genomes, 24 MAGs, 245 unbinned contigs, and 342 reference sequences (**Fig. 3**). Consistent with previous findings [27,29,42,43], our phylogenetic tree revealed that the NosZI is predominantly restricted to the Pseudomonadota, whereas the NosZII showed broader taxonomic distribution across several phyla (**Fig. 3**). Most NosZII sequences affiliate with Bacteroidota, specifically the family *Flavobacteriaceae* (**Fig. 3**). NosZII sequences were also affiliated with other phyla, including Gemmatimonadota, Acidobacteriota and Chloroflexota. While the NosZIII was reported to be associated with diverse taxa [26], this gene was not detected in the isolate genomes or assembled contigs in our dataset, suggesting low prevalence of clade III N_2_O reducers in the studied sandy sediments.

**Figure 3.**
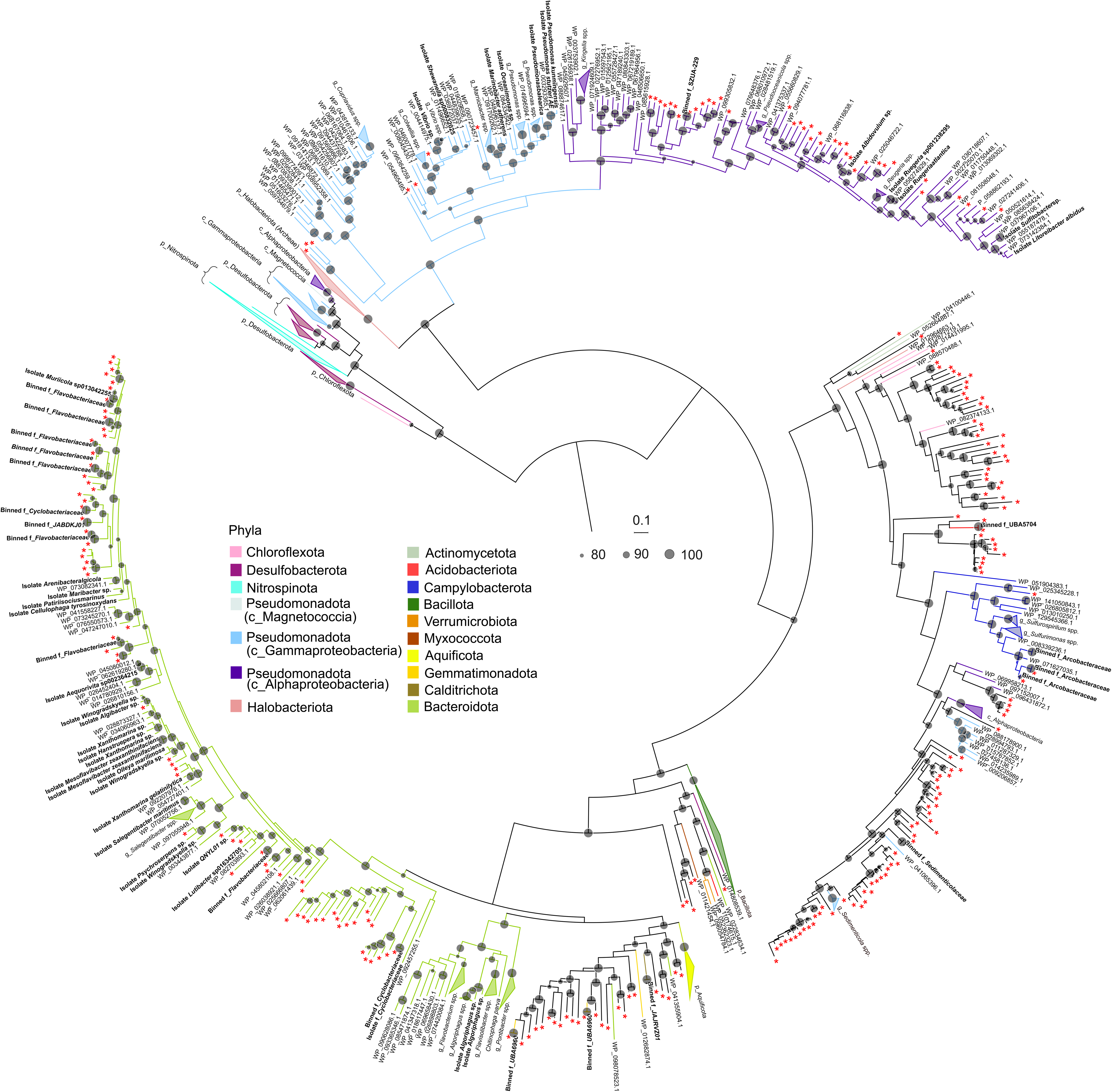
Diversity of NosZ in permeable sediments. The maximum-likelihood phylogenetic tree of NosZ amino acid sequences was generated using Q.pfam+I+G4 substitution model (via IQ-TREE) and was midpoint rooted. The NosZ amino acid sequences derived from MAGs (n = 24) and isolate genomes (n = 36) are shown in **bold**, whereas sequences from unbinned contigs (n = 245) are marked with an asterisk (*). Additionally, 342 reference NosZ sequences were included. Taxonomic assignments are indicated by branch colours on the tree.

### An isolate collection of marine microbes uncovers novel mediators of N_2_O cycling

To complement the metagenomic insights into the potential mediators of N_2_O cycling, we conducted an isolation campaign to obtain high-quality genomes, confirm metabolic activities, and gain ecophysiological insights into key N_2_O cycling microorganisms. To the best of our knowledge, this cultivation effort generated the largest collection of permeable sediment isolates (n = 95) to date. These spanned four main phyla, including Bacillota (n = 6), Actinobacteriota (n = 4), Bacteroidota (n = 36), Alphaproteobacteria (n = 29) and Gammaproteobacteria (n = 20). These isolates represent a considerable fraction of the community’s phylogenetic diversity observed from metagenome-resolved data, including dominant lineages. Based on GTDB, our isolate collection includes two novel bacterial genera and 57 novel bacterial species (**Fig. 2B, Table S4**).

Among these isolates, 12 *Alphaproteobacteria* (*Rhodobacteraceae*, n = 5) and 7 *Gammaproteobacteria* encode a complete set of denitrifying genes, including clade I *nosZ*, indicating capacity to completely reduce NO_3_ ^−^ to N_2_. Another 12 proteobacterial isolates encode denitrification genes except *nosZ*, suggesting they produce N_2_O through partial denitrifications. The partial denitrification phenotype was experimentally validated in two isolates, an *Antarctobacter* sp. and *Sedimentitalea* sp. of the *Rhodobacteraceae* family (Alphaproteobacteria) (**Fig. 4**). The *Antarctobacter* sp. has the genomic capacity to mediate stepwise NO_3_ ^−^ reduction to N_2_O (NO_3_^−^→ NO_2_^−^_→_ NO _→_ N_2_O) via Nar, NirK, and Nor, while *Sedimentitalea* sp. Is predicted to only mediate NO_2_ ^−^ reduction to N_2_O (NO_2_^−^_→_ NO _→_ N_2_O) via NirS and Nor (**Fig. 4B, C**). Consistently, the *Antarctobacter* sp. isolate completely reduced 917 ± 30 µM NO_3_ ^−^ to N_2_O after 24 hours of incubation with the expected 1:1 stoichiometry, after transiently producing small quantities of NO_2_^−^. As expected, the strain also produced N_2_O when supplemented with NO_2_^−^; however, this was a less efficient process and only ∼62% of the initial NO_2_ ^−^ was reduced to N_2_O after 24 hours of incubation, suggesting delayed NO reduction (**Fig. 4B**). Also consistent with the genomic predictions, the *Sedimentitalea* sp. isolate could not consume NO_3_^−^ but did convert NO_2_^−^ to N_2_O after an initial lag (**Fig. 4C**). Given both isolates lacked the N_2_O reductases, they were unable to perform the last step of denitrification and thereby emitted N_2_O as their primary end product.

**Figure 4.**
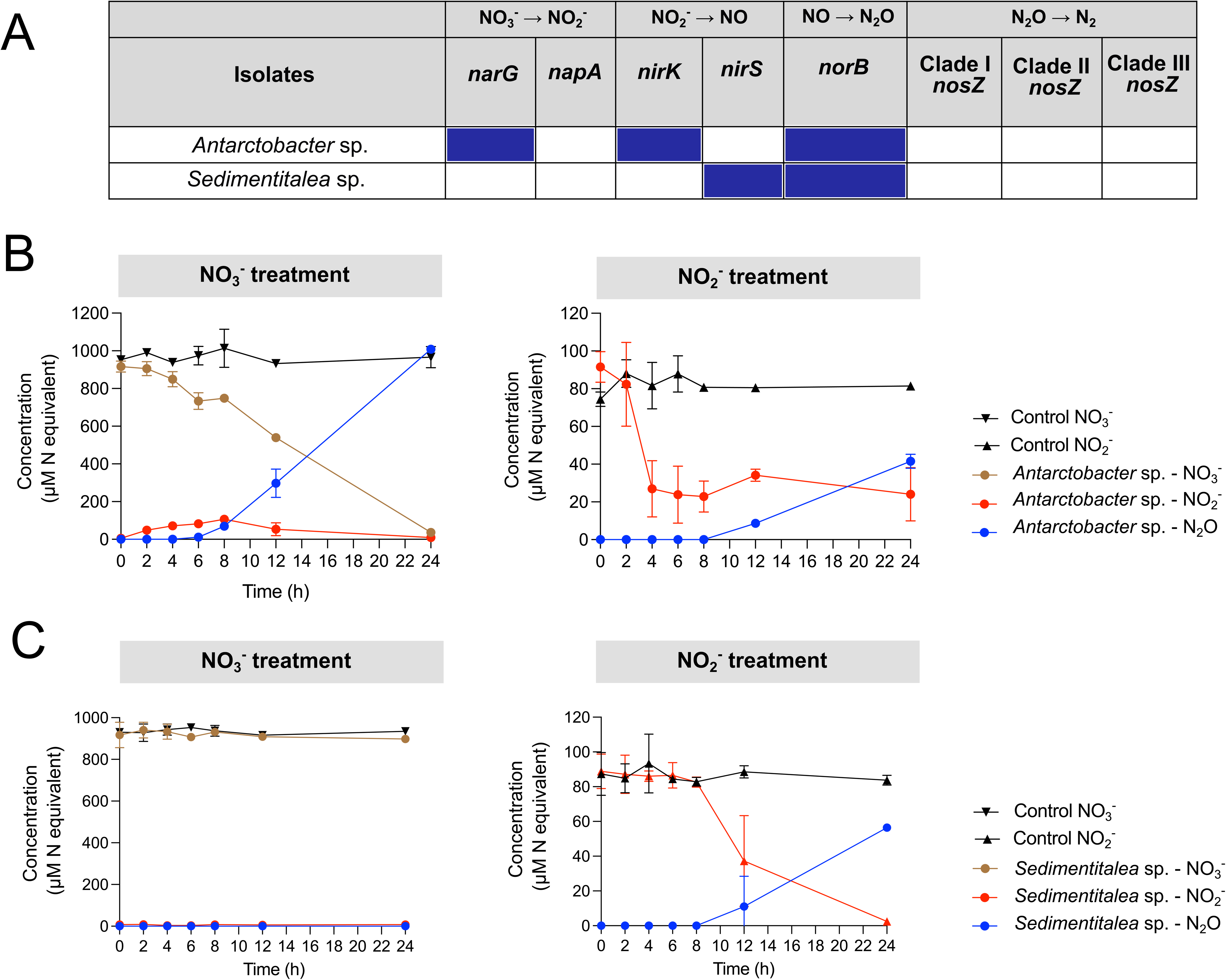
Partial denitrification genotype and phenotype of selected bacterial isolates. **(A)** Presence and absence of denitrifying genes in two isolates. (**B**) Partial denitrification phenotypes in *Antarctobacter* sp. (**C**) Partial denitrification phenotypes in *Sedimentitalea* sp. Both isolates were grown on Difco 2216 Marine Broth with either 1 mM NaNO_3_ for NO_3_^−^ treatment or 0.1 mM NaNO_2_ for NO_2_^−^ treatment under anoxic conditions. Data are presented as mean ± SD, n = 3 biologically independent samples, with media-only vials (n = 2) as negative controls.

On the other hand, most isolates showed widespread capacity for N_2_O reduction. Clade I *nosZ* genes were identified in the 12 Proteobacteria capable of complete denitrification. In contrast, clade II *nosZ* genes were identified in 20 *Flavobacteriaceae* and three *Cyclobacteriaceae* isolates (Bacteroidota). Among these, four encoded the clade II *nosZ* without any denitrifying genes, while 16 co-encoded *nosZ* with *nor* for NO reduction to N_2_O. Notably, some *Flavobacteria*ceae isolates (e.g. *Lutibacter* sp., *Algibacter* sp. and *Winogradskyella* sp.) also encoded *nrf* for DNRA whereas others (e.g. *Arenibacter algicola* or *Xanthomarina* sp.) also encoded the *nirK* genes for NO_2_ ^−^ reduction. Whereas metagenomic predictions indicate that the most dominant *Flavobacteriaceae* lineages within permeable sediments are exclusive N_2_O reducers, results from microbial isolates suggest that *Flavobacteriaceae* members can variably mediate NO_2_ ^−^ reduction, DNRA along with N_2_O reduction.

Further genomic analyses on these isolates revealed their broader metabolic breadth (**Table S4**). Most of them encode multiple terminal oxidases for aerobic respiration while some (e.g. *Lutibacter* sp. or *Oceanimonas* sp.) also encode group 3b or 3d [NiFe]-hydrogenases, which contribute to fermentative H_2_ production within these sediments as previously reported [14]. This suggests that N_2_O-cycling microorganisms can be extremely versatile switching between aerobic respiration, anaerobic respiration and fermentation depending on availability of electron acceptors. Additionally, many isolates encode sulfide-quinone oxidoreductase (Sqr) for sulfide oxidation, while others encoded carbon monoxide dehydrogenase for trace gas oxidation or rhodopsin for phototrophy, suggesting permeable sediment microbes can variably use organic, inorganic and solar energy sources. Altogether, these culture-based insights support our previous genome-resolved metagenomic predictions [38,41] that permeable sediment microorganisms are highly metabolically flexible, possibly as a response to frequent environmental fluctuations.

### Clade II *nosZ* encoding bacteria from permeable sediments show low affinity towards N_2_O

To further determine the physiological characteristics of N_2_O reducers in permeable sediments, we investigated N_2_O consumption by four marine bacterial isolates capable of aerobic organoheterotrophic growth. Two strains, *Olleya marilimosa* and *Winogradskyella* sp. (both *Flavobacteriaceae*), carry the clade II *nosZ* gene but lack key denitrification genes (i.e. *nirS* or *nirK*). In contrast, *Marinobacter adherens (Oleiphilaceae)* and *Oceanimonas* sp. (*Aeromonadaceae)* carry denitrification genes and clade I *nosZ* (**Fig. 5A**). These four isolates grew aerobically in Difco 2216 Marine Broth using yeast extract and peptone as the carbon and energy sources (**Fig.5 A, D,G and J**) and quickly reduced all headspace N_2_O under anoxic conditions (**Fig. 5B, C, D and E**). Whole-cell kinetic measurements revealed that N_2_O reduction followed Michaelis-Menten-type kinetics in all isolates (**Fig. 5B, C, D and E**). The mean apparent half-saturation constants [K_m(app)_] of clade I *nosZ* strains *M. adherens* and *Oceanimonas* sp. were 94.4 µM (95% CI: 47 −230) and 34.1 µM (95% CI: 19 - 72). In contrast, [K_m(app)_] values of clade II *nosZ* flavobacterial isolates *O. marilimosa* and *Winogradskyella* sp. were considerably lower at 10.1 µM (95% CI: 3.9 – 25) and 11.3 µM (95% CI: 5.6 – 22.3), respectively.

**Figure 5.**
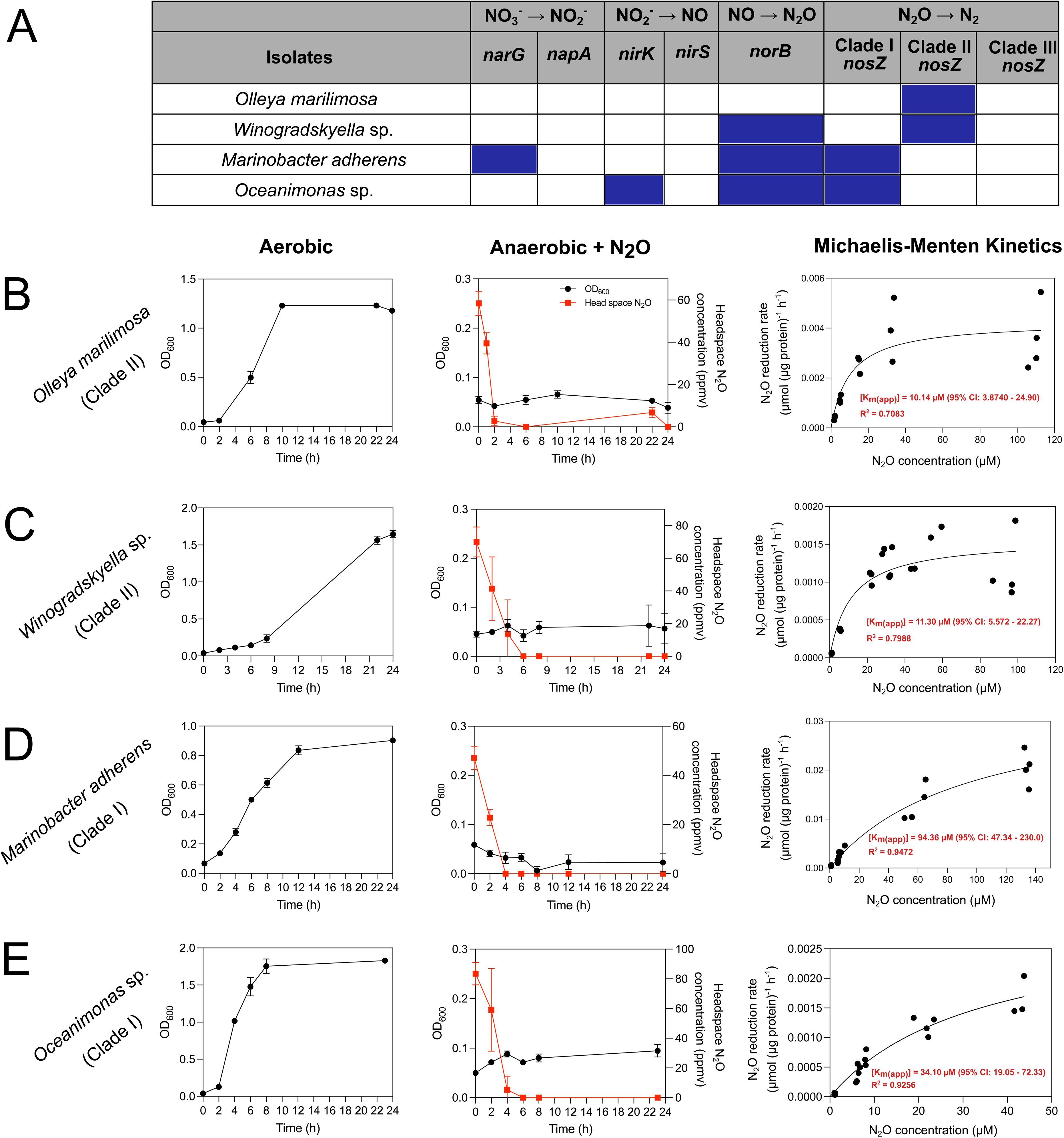
N_2_O reduction genotype and phenotype of clade I and II N_2_O reducing isolates. **(A)** The presence and absence of *nosZ* genes and other denitrifying genes in selected isolates based on whole genome sequencing and metabolic annotation. (**B-D**) Aerobic growth and N_2_O reducing activity of *O. marilimosa* and *Winogradskyella* sp. (clade II *nosZ* isolates) and *M. adherens* and *Oceanimonas* sp. (clade I *nosZ* isolates) respectively. Isolates were grown on Difco 2216 Marine Broth aerobically or anaerobically with N_2_O as the sole terminal electron acceptor. Growth was determined by optical density measurements at 600 nm, followed by measurements of N_2_O concentration in the headspaces of the culture vials. Kinetics of N_2_O reduction of four isolates were calculated based on a Michaelis-Menten non-linear regression model. All experiments were performed in triplicate. Data are presented as mean ± standard deviation (SD).

We subsequently compared the N_2_O sink strength of our permeable sediment isolates with previously characterized N_2_O reducing bacteria from other ecosystems (**Fig. 6**). Clade II *nosZ* flavobacterial isolates in this study had much higher [K_m(app)_] values than most clade II *nosZ* microorganisms reported to date such as *Anaeromyxobacter dehalogens* (1.34 µM) [44] or *Dechloromonas aromatica* RCM (0.324 µM) [45] (**Fig. 6A)**, indicating a relatively low affinity for N_2_O. In addition to substrate affinity, we also compared their ‘catalytic efficiencies’ (V_max_/K_m_) to understand their ability to reduce N_2_O under low N_2_O concentrations (**Fig. 6B**). While the maximum reduction rates (V_max_) of our isolates (except the clade I NosZ containing *M. adherens*) were comparable to the average of the other bacteria, their ‘catalytic efficiencies’ (V_max_/K_m_) was low due to their low apparent affinity towards N_2_O (**Fig. 6B, Table S5**). These results suggest that, among known N_2_O reducing bacteria, our isolates have relatively weak N_2_O sink strength under both high and low substrate conditions regardless of their clade type.

**Figure 6.**
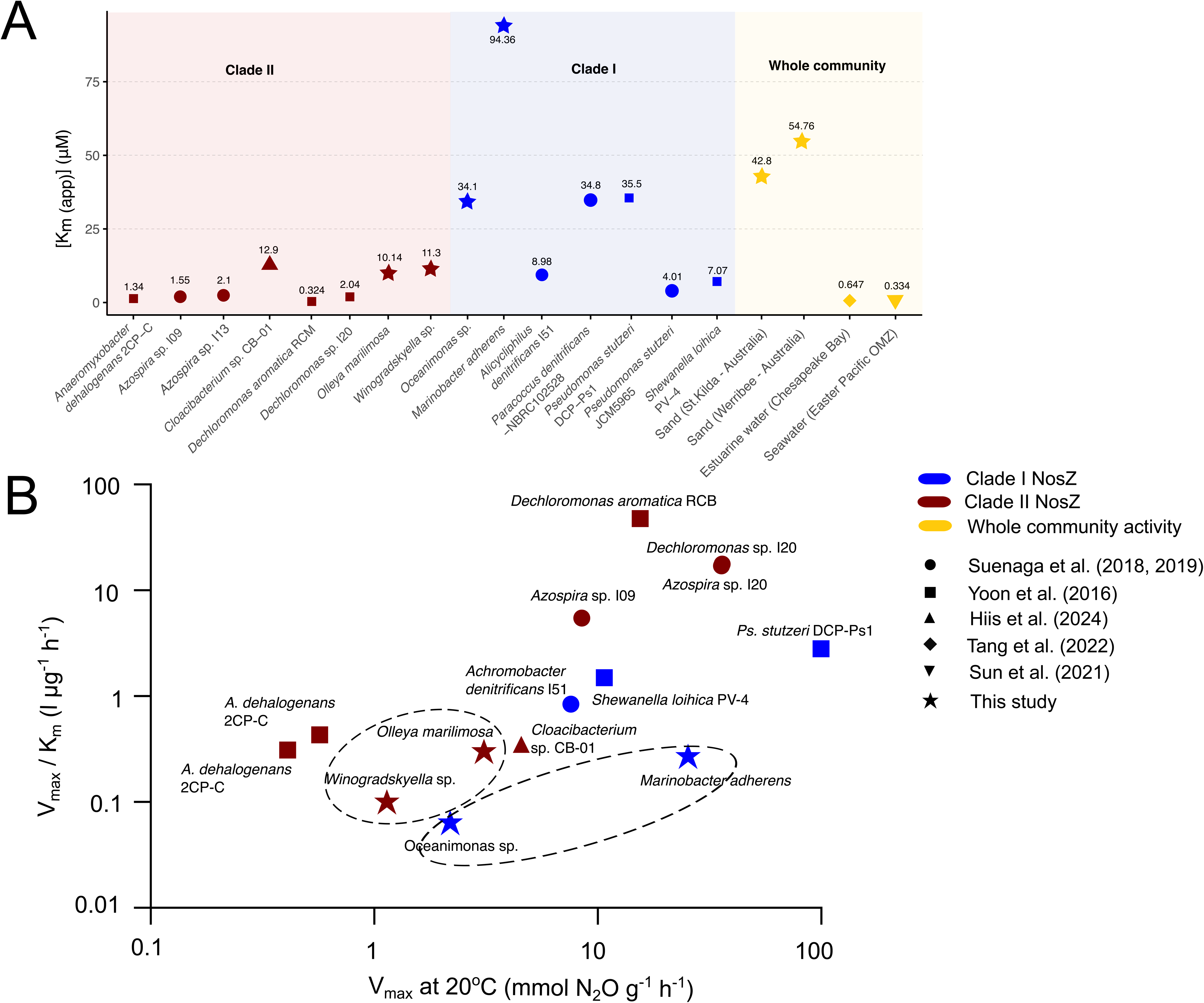
Comparison of N_2_O reduction biokinetics between different N_2_O reducing microorganisms. **(A)** Affinities for N_2_O reported for N_2_O-reducing isolates and whole communities from this and previous studies [40,44–48] (B) N_2_O sink strength of various microorganisms from different ecosystems, determined by plotting the ‘catalytic efficiency’ (V_max_/K_m_) against the maximum reduction rates (V_max_). To enable consistent comparison, we converted all V_max_ values to mmol N_2_O g^−1^ cell dry weight h^−1^, assuming the total protein amounts account for 55% of the total dry biomass [49,50]. We also converted all values to V_max_ at 20°C assuming the rates increase exponentially with temperature by the factor of 2.5 per 10°C increase in temperature (Q_10_ = 2.5), as previously described [46].

## Discussion

Through integrative approaches, we demonstrate that N_2_O cycling in permeable sediment is mediated by a delicate balance between microbial production and consumption. Consistent with previous studies [33,38,41], our results show that permeable sediments host diverse microorganisms with partial denitrification pathways (i.e. lacking one or more genes required for complete denitrification). Some of the most abundant members of the community (e.g. *Rhodobacteraceae* members) possessed either nitrite reductase and/or nitric oxide reductase, but lacked NosZ, thus producing N_2_O as the terminal product of denitrification and contributing to N_2_O emissions. Furthermore, we experimentally confirmed this partial genotype in two *Rhodobacteraceae* isolates (i.e. *Antarctobacter* sp. and *Sedimentitalea* sp.), which produced N_2_O from either NO_3_^−^ or NO_2_^−^ without further reducing it to N_2_, validating metagenomic and culture-based genomic predictions. At the ecosystem level, denitrification is recognized as a collective effort performed by different community members and partial denitrification can be more common across diverse terrestrial and aquatic environments [21,51,52]. Partitioning of denitrification steps among community members could alleviate competition between enzymes acting on the same substrates and enhance community flexibility and resilience under environmental disturbances [51]. Furthermore, specialising in a certain step allows partial denitrifiers to minimise resource allocation to denitrification, enabling broader metabolic flexibility to adapt to complex, dynamic and oligotrophic ecosystems [52]. Given that permeable sediments are highly dynamic with strong spatiotemporal variation in electron donor and acceptor availability, partial denitrification is more prevalent than complete denitrification in these systems. Although we did not directly measure N_2_O production rates in this study, the high [K_m(app)_] values for N_2_O reduction of St. Kilda and Werribee sediments suggest that the N_2_O reducing community in these sites is adapted to high N_2_O concentrations potentially generated by these partial denitrifiers.

Despite substantial capacity for N_2_O production, we provide biogeochemical, metagenomic and culture-based evidence that most N_2_O is consumed by the sediment community before being emitted into the atmosphere. A diverse consortium of N_2_O-reducing microorganisms carrying the clade I and II *nosZ* gene were detected, which efficiently reduce N_2_O to N_2_ before being released into the atmosphere. These microorganisms span several phyla including Bacteroidota, Pseudomonadota, Campylobacterota, and Gemmatimonadota, with metabolically versatile members of the *Flavobacteriaceae* (Bacteroidota) carrying clade II *nosZ* genes particularly dominant. Our results indicate that most flavobacterial isolate genomes and MAGs lack canonical genes required for dissimilatory denitrification, suggesting that these bacteria are unlikely to produce N_2_O via the conventional denitrification pathway. Instead, by expressing the clade II *nosZ* gene, they function as N_2_O respirers that scavenge N_2_O generated by other microorganisms. Moreover, their N_2_O reducing capacity, together with the ability to use diverse energy sources, electron donors, and electron acceptors, reflects their metabolic flexibility that allows them to thrive under frequent oxic-anoxic shifts of these sediments. In this study, clade III *nosZ* was scarcely detected in the top sediment layers. Previous work has suggested that clade III *nosZ* carrying bacteria, such as members of Desulfobacterota or Nitrospinota phyla, are more likely to inhabit anoxic environments [26], whereas the highly dynamic redox conditions of permeable sediments likely limit their occurrence.

In rippled permeable sediments where solutes such as NO_3_^−^ and N_2_O are continually exchanged via advective porewaters [9,53], N_2_O-reducing microorganisms might be expected to rapidly scavenge N_2_O with great sink strength. This hypothesis also stemmed from our observation of few detectable *in situ* N_2_O concentrations, suggesting that N_2_O reducers efficiently consume N_2_O as it is produced, preventing its accumulation in the environment. However, contrary to our hypothesis, our physiological work demonstrates that bacterial isolates from permeable sediments exhibited relatively weak N_2_O-reducing capacity, as reflected in both average maximum reduction rates (V_max_) and low catalytic efficiency (V_max_/K_m_) compared with other known N_2_O-reducing bacteria. This unexpected finding raises questions about the mechanisms regulating N_2_O cycling within these sediments. The relatively weak sink strength of these isolates may likely reflect their adaptation to the physical characteristics of these sediments. Reactive transport models have suggested that the majority of denitrification in permeable sediments is driven by NO_3_^−^ fed from overlying water [54,55]. In these sediments, advective flow and transport dynamics can restrict interactions between oxic and anoxic zones [54]. As a result, under calm conditions without physical disturbances such as intense waves, once NO_3_^−^ enters the reduced reaction zone and is denitrified to N_2_O, long residence time would allow N_2_O reducers to fully reduce N_2_O to N_2_ without the need to invest in high-affinity or high-rate N_2_O reductases. This aligns with previous model predictions showing minimal N_2_O efflux under calm wave conditions which are predominant at these studied sites, correlating with high Damköhler number, a dimensionless ratio of reaction to transport rates [37]. Under more hydrodynamically active conditions, more oxygen is introduced into the sediment, limiting the opportunities for denitrification and N_2_O production to take place. It is only under specific intermediate conditions of sediment flushing that significant amounts of N_2_O escapes the sediment, making it a rare occurrence (<3% of the time) [37].

Altogether, our findings suggest a potential cross-feeding interaction between N_2_O producers and non-denitrifying N_2_O reducers, which efficiently buffers the emissions of this potent greenhouse gas in eutrophic permeable sediments. The efficiency of N_2_O removal by clade II N_2_O-reducing bacteria may be attributed to the unique physical characteristics of these sediments rather than their biokinetic parameters. This suggests that physical characteristics of these sediments not only influence dynamics of greenhouse gases, but also drive the adaptation and ecophysiological strategies of resident microbial communities. Future work should investigate how these microorganisms respond and mediate N_2_O emissions under variable hydrodynamic regimes.

## Materials and Methods

### Sampling and *in situ* N_2_O level measurements

Different sampling campaigns were conducted at St. Kilda Beach (37.863159°S, 144.971026°E) and Werribee Beach (37.58132°S, 144.42112°E) between 2022 and 2024 for different purposes: i) bacterial isolation (02/06/2022, 03/08/2022, 19/10/2022 and 11/01/2023), ii) slurry incubations (10/05/2024 for St. Kilda samples; 23/09/2024 and 20/08/2025 for Werribee samples) and iii) *in situ* N_2_O concentrations (24/05/2024, 05/11/2024 for Werribee and 15/05/2024, 01/10/2024 and 18/11/2024 for St. Kilda). Subtidal surface sediment (0-5 cm deep) was collected in sterile 50 mL Falcon tubes and were immediately transported to the laboratory and stored at 4°C overnight before experiments. The sediments for metagenomic sequencing were also sub-sampled, flash-frozen in liquid nitrogen and stored at −20°C until further analysis. Seawater samples were collected for slurry incubations and were filtered (0.22 µm) to remove most pelagic species. *In situ* N_2_O levels were measured from pore-water samples collected with a piezometer connected with a rubber tube and over-filled in 12 mL exetainers. Three replicates were collected, each spaced 2 m apart. Samples were preserved by adding 50 µL saturated HgCl_2_ and stored at room temperature. For dissolved N_2_O concentration measurement, a 4 mL helium headspace was introduced into each 12.5 mL Exetainer vial, shaken and equilibrated at room temperature before being analysed by gas chromatography described below.

### Slurry incubations

Sediment experiments were conducted in order to determine N_2_O consumption capability of the sediment community. Slurries were prepared with 5 g of wet sediment and 50 mL filtered seawater in 120 mL serum vials before sealing with a butyl rubber stopper (Sigma-Aldrich) and Wheaton closed-top seals (Sigma-Aldrich). Serum vials were purged with helium (He) gas for 10 minutes with vigorous shaking every five minutes to exclude O_2_ and create anoxic conditions. The headspace of vials was amended with N_2_O (Air Liquide Australia Limited) to give initial mixing ratios of approximately 10 (0.8 _µ_M) or 1,000 ppmv (108 _µ_M). Before measurement, 2 mL He was injected into the headspace, then 2 mL of sample was taken from the headspace and injected into a He pre-purged 3 mL exetainer. Samples were analysed using a VICI Trace Gas Analyser with a limit of detection of 0.05 ppmv.

### Metagenomic sequencing and assembly

Environmental DNA was extracted from six sediment samples collected at St.Kilda and Werribee Beach using the PowerSoil® DNA kit (QIAGEN) following the manufacturer’s instructions. Extractions included a sample-free negative control. Samples were sequenced on an Illumina NovaSeqX Plus (2 × 150 bp) at Monash Genomics & Bioinformatics Platform (Monash University). Sequencing yielded an average of 44,021,402 read pairs per sample, with 1,052 read pairs sequenced in the negative control. The metagenomes were processed using the Metaphor workflow [56]. Specifically, the metagenomes were quality filtered with fastp [57] and co-assembled using MEGAHIT v1.2.9 [58] with default settings. Contigs shorter than 1,000 bp were discarded. The assembled contigs were binned with Vamb v4.1.3 [59], MetaBAT v2.12.1 [60] and CONCOCT v1.1.0 [61]. The three bin sets were subsequently refined using DAS Tool v1.1.6 [62], combined with MAGs previously obtained from St. Kilda permeable sediments [38,41] and dereplicated with dRep v3.4.2 [63] using the parameters -comp 50 -con 10. The completeness, contamination and heterogeneity of the bins were assessed using CheckM2 [64]. Quality thresholds for bins were set as per MIMAG standards [65], retaining only medium (completeness >50%, contamination <10%) and high (completeness >90%, contamination <5%) quality bins for further processing, which were termed metagenome-assembled genomes (MAGs). MAG taxonomy was determined using the Genome Taxonomy Database Release R226 [66] via GTDB-Tk v2.3.2 [67]. CoverM [68] with the option ‘genome’ was used to calculate the relative abundance of each MAG in each sample.

### Bacterial isolation and whole genome analysis

Sediment samples for bacterial isolation were collected at St. Kilda Beach in three sampling campaigns (02/06/2022, 03/08/2022 and 11/01/2023) and at Werribee Beach (19/10/2022). Approximately 1 g wet weight of the surface sediment (0-5 cm deep) was dispersed in 9 mL of sterilised 1× phosphate-buffered saline (PBS), vortexed vigorously for three minutes and sonicated for another three minutes. The resulting slurry was serially diluted with 1× PBS and stored in cryovials at −80 °C with 25% glycerol for further isolation if needed. Aliquots of 100 µL undiluted and serially diluted slurry were spread on Marine Agar 2216 (Difco), modified Marine Agar 2216E, and Marine R2A Agar plates (ingredients in **Table S1**) and incubated aerobically at 30 °C for at least one week until most morphologically different colonies appeared on the plates. A series of enrichments were also performed using 1 mM NaNO_3_ or NaNO_2_ as electron acceptors and either 1 mM sodium acetate, 1 mM glucose, 0.002 g/L starch or 0.5 g/L yeast extract as electron donors. Approximately 5 g of sediments were suspended in 50 mL of a bicarbonate-buffered minimal saltwater medium (MSW) (**Table S1**) in 120-mL serum bottles and purged with N_2_ for three minutes to create anoxic conditions. Enrichments were transferred to fresh medium when NO_3_^−^/NO_2_^−^ was all consumed, using 10% inoculum (v/v). After the third transfer, the slurries were serially diluted, isolated by streaking on Marine Agar 2216 or MSW agar and incubated at the same conditions as mentioned above.

Bacterial stocks were preserved at −80 °C in 25% glycerol for long-term storage. For isolate identification, DNA was extracted by heating 30 µL of colony suspension at 95°C for 15 minutes and immediately frozen at −20°C and the 16S rRNA gene was amplified using the universal eubacterial primers 27F (5’-AGA GTT TGA TCC TGG CTC AG-3’) and 806rB (5’-GGA CTA CNV GGG TWT CTA AT-3’) or 1492R (5’-GGT ACC TTG TTA CGA CTT-3’). Purified PCR products were Sanger sequenced (Monash Genomics & Bioinformatics Platform, Victoria, Australia) and the obtained sequences were compared with those available in the GenBank database using BLAST on NCBI (National Center for Biotechnology Information) website.

A total of 95 bacterial isolates from permeable sediments collected in St. Kilda and Werribee beaches were selected for whole genome sequencing. The genomic DNA of isolates was extracted using the Genomic DNA Kit (Meridian Bioscience) following the manufacturer’s instructions. Samples were sequenced on Illumina NovaSeq 6000 platform at GENEWIZ (AZENTA Life Sciences). Raw sequences were quality filtered with the BBTools suite v.38.81 (https://sourceforge.net/projects/bbmap/), using BBDuk to remove the 151^st^ base, trim adapters, filter PhiX reads, trim the 3’ end at a quality threshold of 15 and discard reads below 50 bp in length. Processed short reads were assembled using Unicycler v.0.4.7 [69]. The assembled genome quality was assessed with CheckM2 [64] and taxonomic classification of each isolate was performed using GTDB-tk v2.1.1 [67].

### Metabolic annotation

Open reading frames (ORFs) were predicted using Prodigal v2.6.3 [70] for isolate whole genome and MAGs, then annotated using DIAMOND v2.0.15 [71] for alignment against a custom set of 51 metabolic marker protein databases (https://doi.org/10.26180/c.5230745) covering the major pathways for aerobic and anaerobic respiration, energy conservation from organic and inorganic compounds, carbon fixation, nitrogen fixation and phototrophy. Gene hits were filtered by a minimum percentage identity score by protein: 80% (PsaA), 75% (HbsT), 70% (PsbA, IsoA, AtpA, YgfK and ARO), 60% (CoxL, MmoX, AmoA, NxrA, RbcL, NuoF, [FeFe]-hydrogenases and NiFe Group 4 hydrogenases) or 50% (all other genes). To determine gene abundance in the community, read counts of each gene were normalized to reads per kilobase million (RPKM) and then divided by the mean abundance of 14 universal single-copy ribosomal marker genes (in RPKM, obtained from the SingleM v0.13.2 package) [72].

### Phylogenetic analysis

To understand the distribution and diversity of microorganisms in marine permeable sediments involved in the nitrogen cycling and N_2_O emission, a phylogenomic tree was constructed from 95 bacterial isolate whole genomes and 97 MAGs. Genomes were aligned based on the AR53/BAC120 marker set of GTDB-Tk (v2.4.0) using the “align” option and the tree was generated by maximum likelihood estimation with the ultrafast bootstrap value of 1,000 and model Q.pfam+I+G4 using IQ-TREE [73]. In addition, we also analysed the diversity of clade I and clade II *nosZ* genes present in permeable sediments. To do so, we retrieved the NosZ amino acid sequences derived from unbinned and binned contigs and isolate genomes. Any sequences shorter than 300 amino acids were excluded. We also included the NosZ amino acid sequences from the Greening Lab database (https://doi.org/10.26180/c.5230745) and clade III NosZ sequences from [26] as reference. Sequences were aligned using MAFFT v7.526 [74] and trimmed with TrimAL 1.2rev59 [75] with default settings. Maximum-likelihood trees were constructed with the ultrafast bootstrap value of 1,000 and model Q.pfam+I+G4 using IQ-TREE [73]. The tree was visualised and midpoint-rooted using the Interactive Tree of Life (iTOL) [76].

### Isolate physiological characterisation

The partial denitrification phenotype was tested in two bacterial isolates (i.e. *Antarctobacter* sp. and *Sedimentitalea* sp.). The N_2_O-reducing phenotype was tested in two clade II *nosZ* organisms (i.e. *Olleya marilimosa* and *Winogradskyella* sp.) and two clade I *nosZ* organisms (i.e., *Marinobacter adherens* and *Oceanimonas* sp.). All cultures were grown overnight in Difco 2216 Marine Broth medium under oxic conditions. Overnight cultures were transferred to 50 mL of fresh media so that the starting optical density (OD_600_) reached 0.03. Oxic treatment remained as unamended air while anoxic treatments were purged with He to exclude O_2_ as described above. For partial denitrification phenotype analysis, cultures were amended with either 1mM NaNO_3_ or 0.1 mM NaNO_2_ and the concentrations of NO_x_ (NO ^−^ + NO_2_ ^−^) and NO_2_ ^−^ were determined spectrophotometrically on a Lachat Quickchem 8000 following the procedures in Standard Methods for Water and Wastewater [77]. Headspace gas (2 mL) was sampled into He purged 12-mL exetainers to monitor N_2_O production. For N_2_O reduction phenotype analysis, the headspace of anoxic vials were amended with N_2_O as the main electron acceptor to give initial mixing ratios of approximately 100 and 1,000 ppmv. Vials were placed on an orbital shaker (150 rpm) at 30°C for 24 hours. To monitor N_2_O concentration changes, 2 mL of headspace gas was sampled into He purged 3-mL exetainers.

### Kinetic analysis

Rates of N_2_O consumption by *O. marilimosa, Winogradskyella* sp., *M. adherens* and *Oceanimonas* sp. and by whole sediment communities from the two studied sites were measured. For axenic cultures, overnight-grown cells under oxic conditions (OD_600_ of 3.0 to 4.0) were inoculated into 50 mL fresh Difco Marine Broth medium in 120-mL serum vials to reach an initial OD_6oo_ of 0.03 then vials were sealed with butyl rubber stoppers. Similarly, slurry was prepared with 5 g of wet sediment and 50 mL filtered seawater in 120-mL serum vials and sealed with butyl rubber stoppers. Vials were purged with He gas to exclude O_2_ as described above. Triplicate cultures or slurries were incubated with different N_2_O concentrations in the headspace ranging from 5 to 1,000 ppmv. Headspace N_2_O concentrations were quantified by gas chromatography at different time intervals. The final N_2_O concentration in the liquid phase was corrected and calculated as previously described [78]. Reaction rates of N_2_O consumption were determined from the slopes of linear regression lines fitted to N_2_O concentration measurements over time. Michaelis-Menten curves and parameters were estimated using the non-linear fit (Michaelis-Menten, least squares regression) function in GraphPad Prism v10.3.1.

## Data availability

All sequences generated from this work were deposited to the NCBI Sequence Read Archive under the BioProject accession numbers PRJNA1347951 for metagenomes and metagenome-assembled genomes and PRJNA1348900 for isolate whole genomes. Bacterial and archaeal MAGs generated from the same sites in previous studies [38,41] were downloaded from NCBI Sequence Read Archive with BioProject accession number PRJNA623061 and PRJNA609151.

## Conflict of interest

The authors declare no conflict of interest.

## Acknowledgement

This work was supported by an ARC Discovery Project (DP210101595; awarded to P.M.L.C, C.G. and W.W.W), an ARC Future Fellowship (FT240100502; awarded to C.G), ARC Discovery Early Career Awards (DE250101210 to P.M.L., DE230100542 to R.L.), Monash International Tuition Scholarships and Monash Graduate Scholarship (awarded to T.N.D).

## Author contributions

C.G, P.L.M.C and W.W.W conceived and supervised the study. Experimental planning and design were conducted by T.N.D, C.G., W.W.W. and P.L.M.C. T.N.D and L.J were responsible for bacterial isolation and culturing. T.N.D, H.P., and W.W.W. were responsible for environmental microcosms and monitoring. T.N.D. and T.F.H were responsible for physiological and kinetic characterization of isolates. Chemical analyses were performed by V.E. Metagenomic and whole genome analyses were performed by T.N.D, F.R., R.L and P.M.L. Resources, supervision and funding were contributed by C.G., P.L.M.C and W.W.W. The manuscript was written by T.N.D and F.R. with contributions from all authors.

## Supplementary materials

**Figure S1** | *In situ* N_2_O concentrations in permeable sediments

**Table S1** | Composition of different media used for isolation of bacteria from permeable sediments in St. Kilda Beach and Werribee.

**Table S2 (xlsx).** Sequencing statistics and abundance of metabolic marker genes in the short read metagenomic data. This includes the calculated average copies per organism for each marker gene, average copies per organism for nitrogen metabolising genes and a list of short read hits and their corresponding match in the database.

**Table S3 (xlsx).** Summary and annotation of dereplicated metagenome-assembled-genomes (MAGs) obtained from this study and previous [38,41] studies. This includes taxonomic information for each MAG, relative abundance of each MAG per sample calculated by CoverM, a summary of metabolic marker genes identified in each MAG, a list of clade II *nosZ* containing MAGs, a list of high-quality MAGs and a full list of metabolic marker genes identified in the MAGs with alignment information and protein sequences.

**Table S4 (xlsx).** Summary of bacterial isolates. This includes taxonomic information for each isolate and a summary of metabolic marker genes identified in each isolate genome.

**Table S5.** Comparison of biokinetic parameters for marine permeable sediment isolates and other N_2_O-reducing bacteria.

**Table S6 (xlsx).** Sequences of metabolic genes in unbinned contigs derived from homology-based searches.

**Table S7 (xlsx).** Measurements of N_2_O concentrations and kinetic analyses. This includes measurement of N_2_O concentrations in slurries and four bacterial isolates.

**Table S8 (xlsx).** Measurements of NO_x_ and N_2_O in two *Rhodobacteraceae* isolates (i.e. *Antarctobacter* sp. and *Sedimentitalea* sp.) with partial denitrification pathways.

**Table S9 (xlsx).** *In situ* N_2_O concentrations in porewaters of St. Kilda and Werribee permeable sediments.

## Notes

### Competing Interest Statement

The authors have declared no competing interest.

### Summary of Updates

1. Author affiliations updated 2. Changed "Figure 1A" on line 1345 to "Fig. 1A" 3. Moved "Figure 5A" from line 288 to line 292 and changed to "Fig. 5A" 4. The figure label in the manuscript on line 293 - "Fig.4 A, D, G, and J" and on line 294 - "Fig. 4B, E, H and K" changed to "Fig. 5B, C, D, and E". 5. The figure label on line 296 changed from "Fig. 5C, F, I and L" to "Fig. 5B, C, D and E" 6. Changed "Figure 6B" on line 314 to "Fig. 6B"

