## Supplementary information for "Flavobacteria buffer nitrous oxide emissions from partial denitrifiers in coastal sediments"

**Table S2 (xlsx).** Sequencing statistics and abundance of metabolic marker genes in the short read meganomic. This includes the calculated average copies per organism for each marker gene, average copies per organism for nitrogen metabolising genes and a list of short read hits and their corresponding match in the database.

**Table S3 (xlsx).** Summary and annotation of dereplicated metagenome-assembled-genomes (MAGs) obtained from this study and [1,2] studies. This includes taxonomic information for each MAG, relative abundance of each MAG per sample calculated by CoverM, a summary of metabolic marker genes identified in each MAG, a list of clade II *nosZ* containing MAGs, a list of high-quality MAGs and a full list of metabolic marker genes identified in the MAGs with alignment information and protein sequences.

**Table S4 (xlsx).** Summary of bacterial isolates. This includes taxonomic information for each isolate and a summary of metabolic marker genes identified in each isolate genome.

**Table S6 (xlsx)**. Sequences of metabolic genes in unbinned contigs based on homology-based searches.

**Table S7 (xlsx).** Measurements of N_2_O concentrations and kinetic analyses. This includes measurement of N_2_O concentrations in slurries and four bacterial isolates.

**Table S8 (xlsx).** Measurements of NO_x_ and N_2_O in two *Rhodobacteraceae* isolates (i.e. *Antarctobacter* sp. and *Sedimentitalea* sp.) with partial denitrification pathways.

**Table S9 (xlsx).** In situ N_2_O concentrations in porewaters of St. Kilda and Werribee permeable sediments.

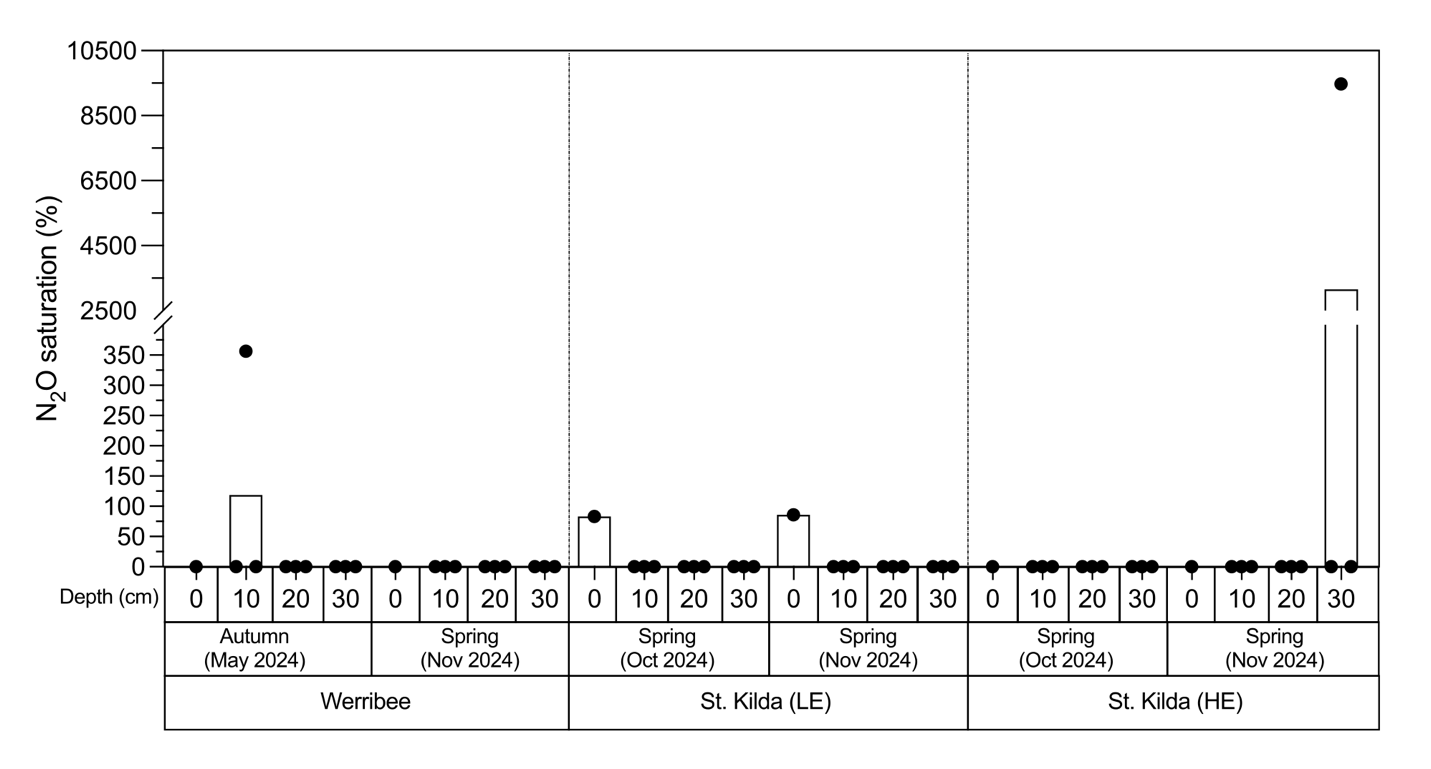

**Figure S1. *In situ*** **N_2_O concentrations in permeable sediments.** Pore-water (0, 10 and 30 cm) concentrations of N_2_O collected at Werribee ( 24/05/2024 and 05/11/2024) and St. Kilda beaches at low energy site (LE) and high energy site (HE) (15/05/2024, 01/10/2024 and 18/11/2024). Pore-water samples were head-spaced with helium gas and analysed using VICI Trace Gas Analyser with the detection limit of 0.05 ppmv.

**Table S1.** Composition of different media used for isolation of bacteria from permeable sediments in Saint Kilda Beach and Werribee.

| **Medium** | **Composition** | **Reference** |
| --- | --- | --- |
| Difco 2216 marine broth | 5g peptone, 1g yeast extract, 0.1g ferric citrate, 19.45g NaCl, 12.6g MgCl_2_.6H_2_O, 6.63 MgSO_4_.7H_2_O, 2.38 g CaCl_2_.2H_2_O, 0.55g KCl, 0.16g NaHCO_3_, 100ml trace element and 1000 mL MQ water. | <https://legacy.bd.com/europe/regulatory/Assets/IFU/Difco_BBL/212185.pdf> |
| Marine agar 2216E | 1g yeast extract, 5g peptone, 0.1 g ferric citrate, 15 g agar, 1000 mL aged sea water | [3] |
| Marine R2A agar | 0.5 g yeast extract, 0.5 g peptone, 0.5.g casein, 0.5 g glucose, 0.5 g soluble starch, 0.5 g sodium pyruvate, 15 agar, 750 mL aged sea water and 250 mL MQ water. | [3] |
| Minimal saltwater medium (MSW) | 20g NaCl, 0.25g NH_4_Cl, 0.2g KH_2_PO_4_, 0.5g KCl, 3g MgCl_2_.6H_2_O, 0.15g CaCl_2_.2H_2_O, 30mL NaHCO_3_, trace element solution 1mL, 0.5 mL Vitamin B_12,_ 1mL vitamin mix, 1mL thiamine, 2mL selenite and tungstate, and 1000mL MQ water. | [4] |

**Table S5.** Comparison of biokinetic parameters for marine permeable sediment isolates and other N_2_O reducing bacteria

| **Organism** | **Clade** | **Temp** | **V_max_** | **V_max20_** | **Km** | **Ref** |
| --- | --- | --- | --- | --- | --- | --- |
|  |  | ^o^C | mol N_2_O g^-1^ DW h^-1^ | mmol N2O g^-1^ h^-1^ at 20^o^C | uM |  |
| *Dechloromonas* sp. I20 (ISO_OTU 07) | II | 30 | 0.090 | 36.0 | 2.04 | [5] |
| *Azospira* sp. I09 | II | 30 | 0.021 | 8.5 | 1.55 |  |
| *Azospira* sp. I13 | II | 30 | 0.090 | 35.8 | 2.1 |  |
| *Dechloromonas aromatica* RCB | II | 30 | 0.039 | 15.5 | 0.324 | [6] |
| *Anaeromyxobacter dehalogenans* 2CP-C | II | 30 | 0.001 | 0.57 | 1.34 |  |
| *Cloacibacterium* sp. | II | 23 | 0.006 | 4.56 | 12.9 | [7] |
| ***Winogradskyella* sp.** | **II** | **30** | **0.0029** | **1.14** | **11.3** | **This study** |
| ***Olleya marilimosa*** | **II** | **30** | **0.0077** | **3.10** | **10.14** |  |
| *Alicycliphilus denitrificans* I51 | I | 30 | 0.019 | 7.6 | 8.98 | [5] |
| *Shewanella Loihica* PV-4 | I | 30 | 0.027 | 10.7 | 7.07 | [6] |
| *Pseudomonas stutzeri* DCP-Ps1 | I | 30 | 0.250 | 99.8 | 35.5 |  |
| *Paracoccus denitrificans* NBRC102528 | I | 30 | 0.003 | 1 | 34.8 | [8] |
| *Pseudomonas stutzeri* JCM5965 | I | 30 | 0.008 | 3.3 | 4.01 |  |
| ***Marinobacter adherens*** | **I** | **30** | **0.064** | **25.4** | **94.36** | **This study** |
| ***Oceanimonas* sp** | **I** | **30** | **0.0055** | **2.19** | **34.1** |  |
